# Sharp-wave ripple associated activity in the medial prefrontal cortex supports spatial rule switching

**DOI:** 10.1101/2022.11.03.515023

**Authors:** Hanna den Bakker, Marie Van Dijck, Jyh-Jang Sun, Fabian Kloosterman

## Abstract

Previous studies have highlighted an important role for hippocampal sharp-wave ripples in spatial alternation learning, as well as modulating activity in the medial prefrontal cortex (mPFC). However, no study so far has investigated the direct influence of hippocampal sharp-wave ripples on mPFC activity during spatial alternation learning. Long Evans rats were trained on a three-arm radial maze to perform a sequence of alternations. Three different alternation sequences needed to be learnt, and while learning a new sequence, the activity in the mPFC was inhibited either directly following sharp-wave ripples in the hippocampus (on-time condition) or with a randomized delay (delayed condition). In the on-time condition the behavioral performance was significantly worse compared to the same animals in the delayed inhibition condition, as measured by a lower correct alternation performance and more perseverative behavior. This indicates that the activity in the mPFC directly following hippocampal sharp-wave ripples is necessary for spatial rule switching.

## INTRODUCTION

Successfully navigating through an environment and learning spatial rules requires cognitive functions, such as memory of rewarded locations, and executive functions while forming behavioral strategies. Both the hippocampus and the medial prefrontal cortex (mPFC) play key roles in these processes. The hippocampus is thought to be important in working memory and memory consolidation, and internalizing a mental map of the environment^1^. When required to form a new spatial strategy due to changing reward contingencies, the mPFC is thought to guide behavior by suppressing what is already known and redirecting attention to newly relevant cues^2^.

Previous studies have shown that inactivation of the hippocampus through chemical lesions causes a behavioral deficit in a spatial alternation task, where the animals must learn to alternate between two arms on a maze^3,4^. Inactivation of the mPFC has also been shown to have an effect on delayed alternation tasks, where it led to a working memory deficit^3,5,6^, and increased perseverative behavior^5^. Attention deficits^7^, premature responses^8^, and decreased VTE behavior^9^ have also been reported as a result of mPFC inactivation. Interestingly, post-training inactivation of the mPFC did not affect behavioral performance^10^, indicating that the behavioral effect seen may be due to problems in guiding attention and selecting behavioral strategies rather than a problem with memory consolidation. In support of this, it was also found that mPFC inactivation selectively impaired the learning of a rule reversal^11,12^, in which the animal is required to suppress learned information and adopt a novel spatial strategy or rule instead.

Emerging evidence further suggests that the interaction between the hippocampus and mPFC plays a key role in the acquisition of new tasks and long-term memory. Lesions aimed at severing the largely ipsilateral functional connections between the hippocampus and mPFC demonstrated that the disruption of their interactions impairs working memory on a spatial alternation task^13^ and spatial reversal learning^14^. Moreover, optogenetic inactivation of hippocampal terminals in the mPFC impaired encoding of rule learning^15^ and blocked novelty-enhanced learning^16^.

Hippocampal-mPFC interactions that are instrumental in learning processes may occur at various times during behavior, such as approach to a choice point^17^, when receiving rewarding feedback^18^ and sleep^19^. In particular, hippocampal sharp-wave ripples (SWRs) – transient high frequency oscillations that are associated with neural ensemble reactivation^20–23^ and memory consolidation^24,25^– form a candidate neural basis for the interaction with the mPFC. Disruption of hippocampal activity specifically during SWRs slowed the acquisition of a spatial alternation^26^, whereas artificial prolongation of hippocampal SWRs using optogenetic excitation improved performance in the same task^27^. SWR manipulation in other working memory tasks showed that behavioral performance is also impaired when disrupting SWRs in a post-training consolidation period^28^, and that this effect is specific to the learnt environment and does not generalize to other environments^29,30^. While SWRs originate in the hippocampus, they are thought to be part of a broader hippocampal-cortical dialogue^1^. Neural activity in the mPFC is modulated by the occurrence of SWRs in the hippocampus^19,21,31,32^, where some neurons increase, and others decrease their firing rate in the 200 ms after SWR onset^31^. It was further found that mPFC modulation is stronger for awake than for sleep hippocampal SWRs^32^, and that a subset of mPFC neurons is selectively modulated by the content of the SWR, i.e. which maze arm is present in the replayed trajectory^21^. These data highlight the potential importance of mPFC activity following SWRs in the hippocampus, although direct causal evidence for this has not previously been reported.

Collectively, these findings point to some intriguing hypotheses about the specific timing of the interaction between the hippocampus and mPFC in spatial learning. Although previous studies have attempted to unravel the functional interaction by chemical and optogenetic manipulations, no study so far has looked at the exact time at which this interaction occurs with respect to neural signatures. The question remains whether the specific timing of hippocampal SWRs, related to the mPFC activity, is the key element that drives behavior in spatial learning.

## RESULTS

### SWR-triggered inhibition of mPFC activity in a spatial alternation task

Adult Long-Evans rats (N=11) were trained across multiple learning blocks over 1-3 days (up to eight 15-minute learning blocks per day) on a spatial alternation paradigm in a symmetrical three-arm radial maze. One arm was randomly selected as central arm and animals had to learn to alternate between the other two arms (left and right arms) while returning to the central arm in between (see Methods and Fig. 1a). Animals reached the initial learning criterium of 80% correct in 12 learning blocks on average (range: 7 to 19 learning blocks). After reaching the learning criterium, the reward contingencies on the maze changed (rule switch) and the animals had to deduct through trial and error the new location of the central, left, and right arms (Fig. 1a). In most animals, two rule switches were performed such that all three arms of the maze served as central arm exactly once (Supplementary Table S1).

**Figure 1.**
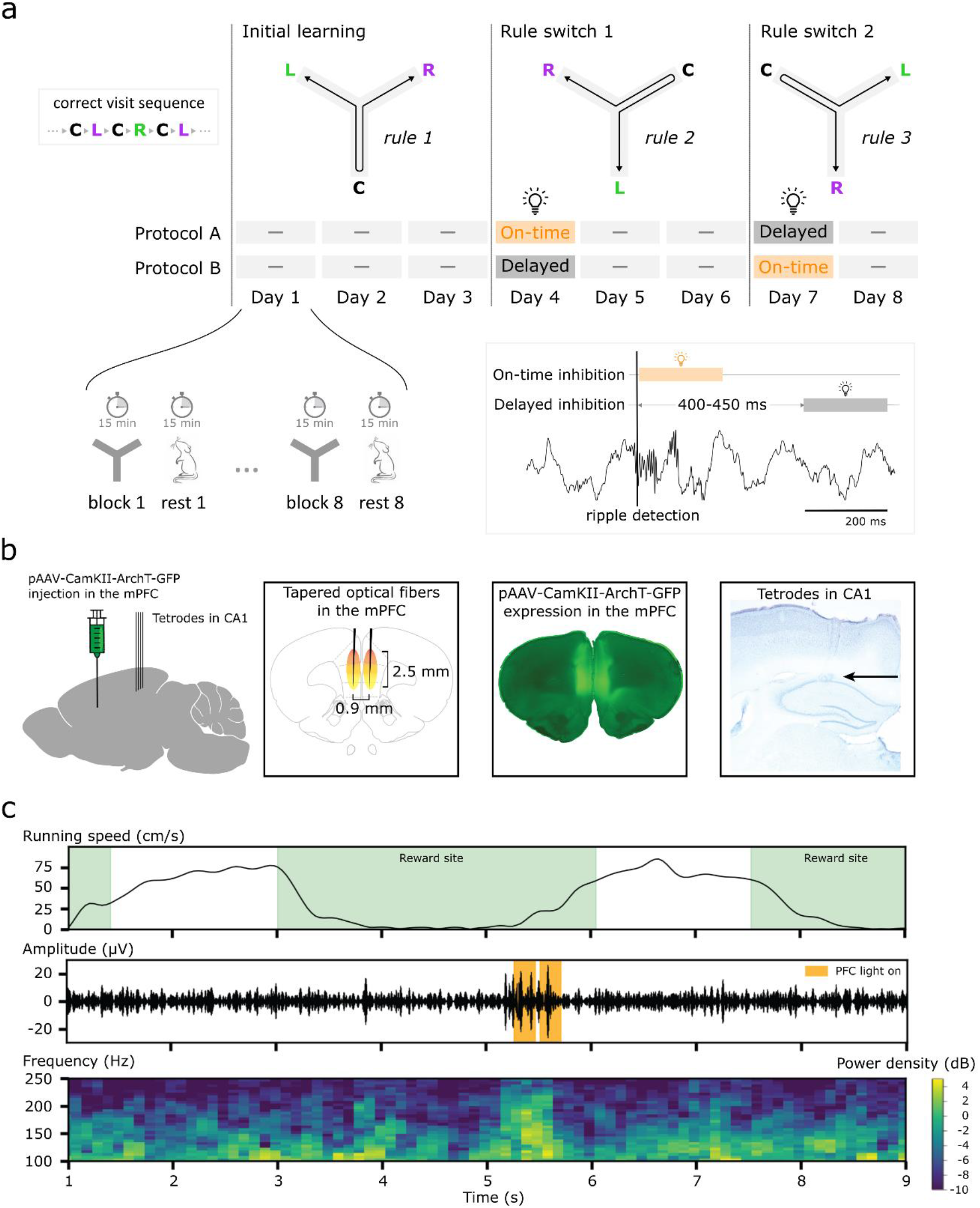
Animals perform a spatial alternation task with SWR-triggered mPFC inhibition. (**a**) Animals had to learn to alternate between left and right arms and visit the center arm in between each alternation. The configuration of the central, left and right arms changed with every rule switch. Each day, animals were placed on the maze where they were allowed to explore freely for 15 minutes, followed by 15 minutes in a dark resting box, until a maximum of 8 learning blocks per day. On experimental day 4 and 7, mPFC activity was inhibited either directly following a SWR detection in the hippocampus (on-time condition) or after a 400-450 millisecond delay (delayed condition). (**b**) pAAV-CamKII-ArchT-GFP was injected into the mPFC and tapered optical fibers were implanted for optogenetic inhibition. Tetrodes were implanted in the hippocampus for SWR detection. (**c**) Example segment of data with the animal’s running speed (top panel), ripple-filtered hippocampal LFP (middle panel), and spectrogram of the hippocampal signal (bottom panel).

Previously, it was shown that activity in the mPFC supports adaptation to rule changes^11,12,14,33^, and that hippocampal SWRs support acquisition of the spatial alternation task^26,27^. Here, we asked if mPFC activity specifically during and immediately following hippocampal SWRs is required for rule switches in the alternation task. On the first day of a rule switch, SWRs in the hippocampus were detected online and activity in the mPFC was inhibited optogenetically either directly following the detection (on-time inhibition), or with a randomized delay of 400-450 milliseconds (delayed inhibition; see Methods and Fig. 1a-b). On days two and three of a rule switch, animals were allowed to learn the new rule to criterium without mPFC inhibition (Fig. 1a).SWRs occurred mostly at the reward sites (overall ripple rate in Hz: M=0.35, SEM=0.05, percentage occurring at reward sites: 75%; example see Fig. 1c), and in the rest periods in between learning blocks (ripple rate during rest periods in Hz: M=0.51, SEM=0.05).

### SWR-triggered inhibition of the mPFC disrupts spatial alternation learning

If SWR-associated activity in the mPFC is required for animals to successfully adapt to the rule switch, we would predict that task performance is negatively affected by on-time inhibition but not delayed inhibition. In support of this hypothesis, we observed that individual animals readily adapted to the changed rule in the delayed inhibition condition but adapted more slowly or not at all in the on-time inhibition condition (Fig. 2). We first quantified the percentage of correct (rewarded) arm visits across all learning blocks on day one of the rule switch (Fig. 2a-b). On average, in the delayed inhibition condition, the percentage of correct visits gradually increased over the eight learning blocks to reach pre-switch levels (Fig. 2b, left). In contrast, in the on-time inhibition condition, the fraction correct visits was near chance for most of the learning blocks and did not reach pre-switch levels. Across all learning blocks, the percentage of correct visits was significantly higher in the delayed versus the on-time inhibition condition (Fig. 2b, right; fraction correct, delayed inhibition: M=0.72, SEM=0.04; on-time inhibition: M=0.54, SEM=0.05; paired samples t-test: t(10)=2.74, *p*=.021). The total number of arm visits was lower in the on-time (M=53.66, SEM=11.53) compared to the delayed (M=70.94, SEM=7.96) inhibition condition, although this effect was not significant (paired samples t-test: t(10)=1.41, *p*=.190).

**Figure 2.**
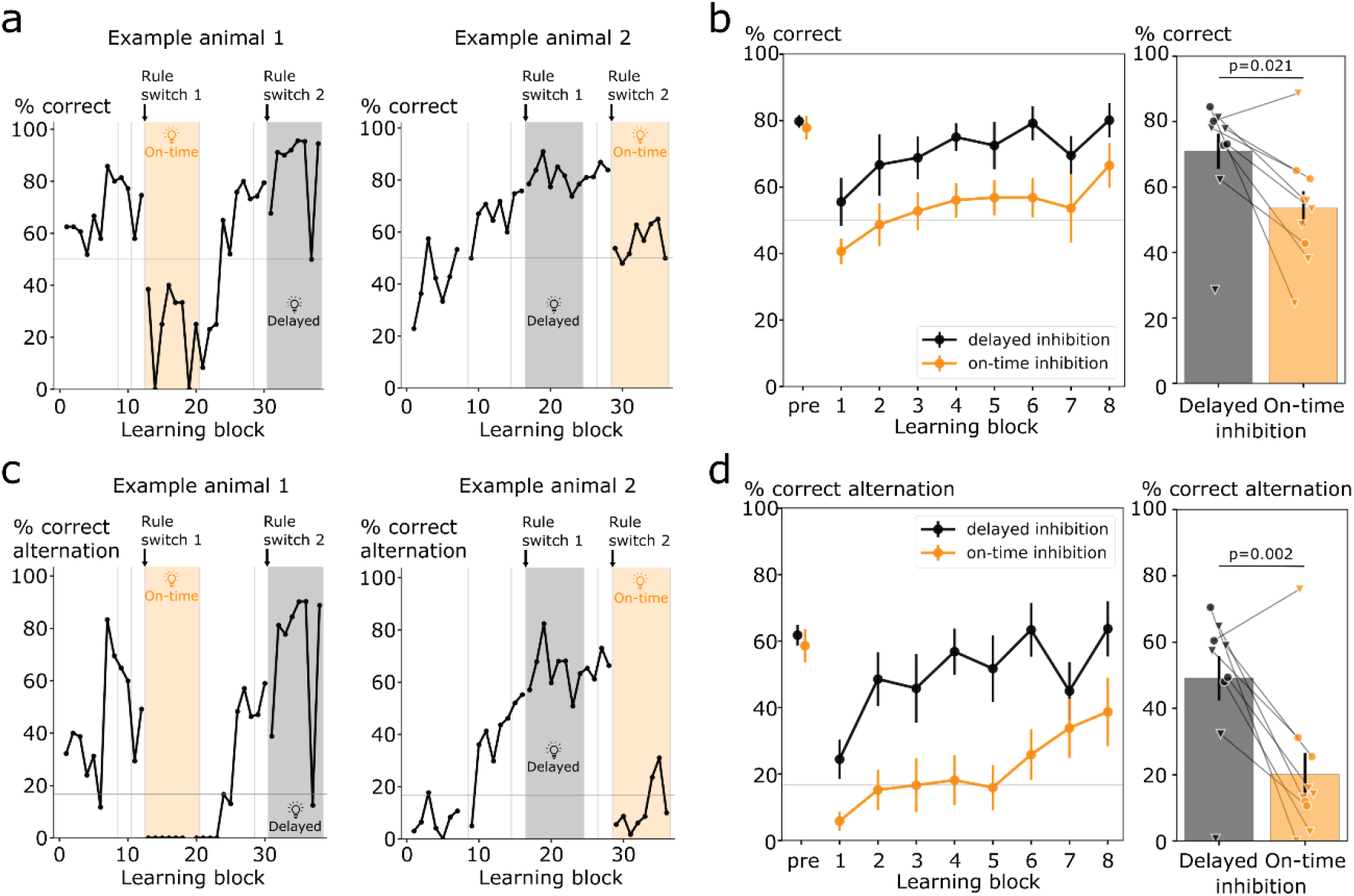
SWR- triggered inhibition of the mPFC disrupts spatial alternation learning. (**a**) Example learning curves of two animals. During the rule switch sessions, mPFC activity was inhibited either immediately following SWRs in the hippocampus (on-time condition) or with a 400-450 delay (delayed condition). (**b**) Left panel: the percentage of correct trials for each 15-minute learning block. ‘pre’ indicates the performance on the previous day. Right panel: the percentage of correct trials over all 15-minute learning blocks between the delayed and on-time conditions. Triangles indicate a rule switch 1 and circles indicate a rule switch 2. (**c**) Example learning curves of the same two animals but expressed in the percentage of correct alternation sequences (e.g., center-left-center-right). (**d**) The percentage of correct alternation sequences in the same configuration as b). The grey line indicates chance level.

The percentage of correct visits is based on binary measure of performance in single visits and provides little insight into the behavioral response patterns in the alternation task. To characterize the behavioral responses in more detail, we analyzed all sequences of 4 visits and computed the fraction of 4-sequences that were consistent with the desired alternation pattern (e.g., center-left-center-right; see Supplementary Fig. S1b). Of the 24 possible 4-sequences, six follow the alternation pattern giving a chance level of 0.16 (Supplementary Fig. S1a, left). We observed that in delayed inhibition conditions animals quickly followed the new alternation sequence and reached pre-switch levels at the end of the day (Fig. 2c-d). In on-time inhibition conditions, animals performed below or at chance level in the first 5 learning blocks and subsequently improved towards the end of the day (Fig. 2d, left). The average alternation performance across all learning blocks was significantly higher in the delayed inhibition condition (fraction of correct alternation sequences, M=0.51, SEM=0.06) than in the on-time inhibition condition (M=0.19, SEM=0.06; Fig. 2d, right; paired samples t-test: t(10)=4.11, *p*=.002).

### On-time mPFC inhibition leads to perseverative behavior

Perseverative behavior can be defined as animals persisting in making the same erroneous choices that are not leading to the most rewarding outcomes. Perseverative behavior has been reported previously as a result of mPFC inhibition^5^. Detailed analysis of the 4 visit sequences allowed us to identify three incorrect behavioral patterns in addition to the correct alternation pattern: running back-and-forth between two arms, visiting all three arms in sequence (circling) and incorrect alternation according to a rule that is not currently in effect (see Supplementary Fig. S1).

Animals showed individual biases for specific error patterns (Fig. 3a). On average, animals engaged in a low level of back-and-forth behavior before the rule switch, which remained largely constant in the learning blocks following the rule switch (Fig. 3b). There was no significant difference in back-and-forth behavior between the on-time and delayed inhibition conditions (Fig. 3b; delayed inhibition: M=0.17, SEM=0.01; on-time inhibition: M=0.15, SEM=0.03; paired samples t-test: t(10)=0.65, *p*=.531).

**Figure 3.**
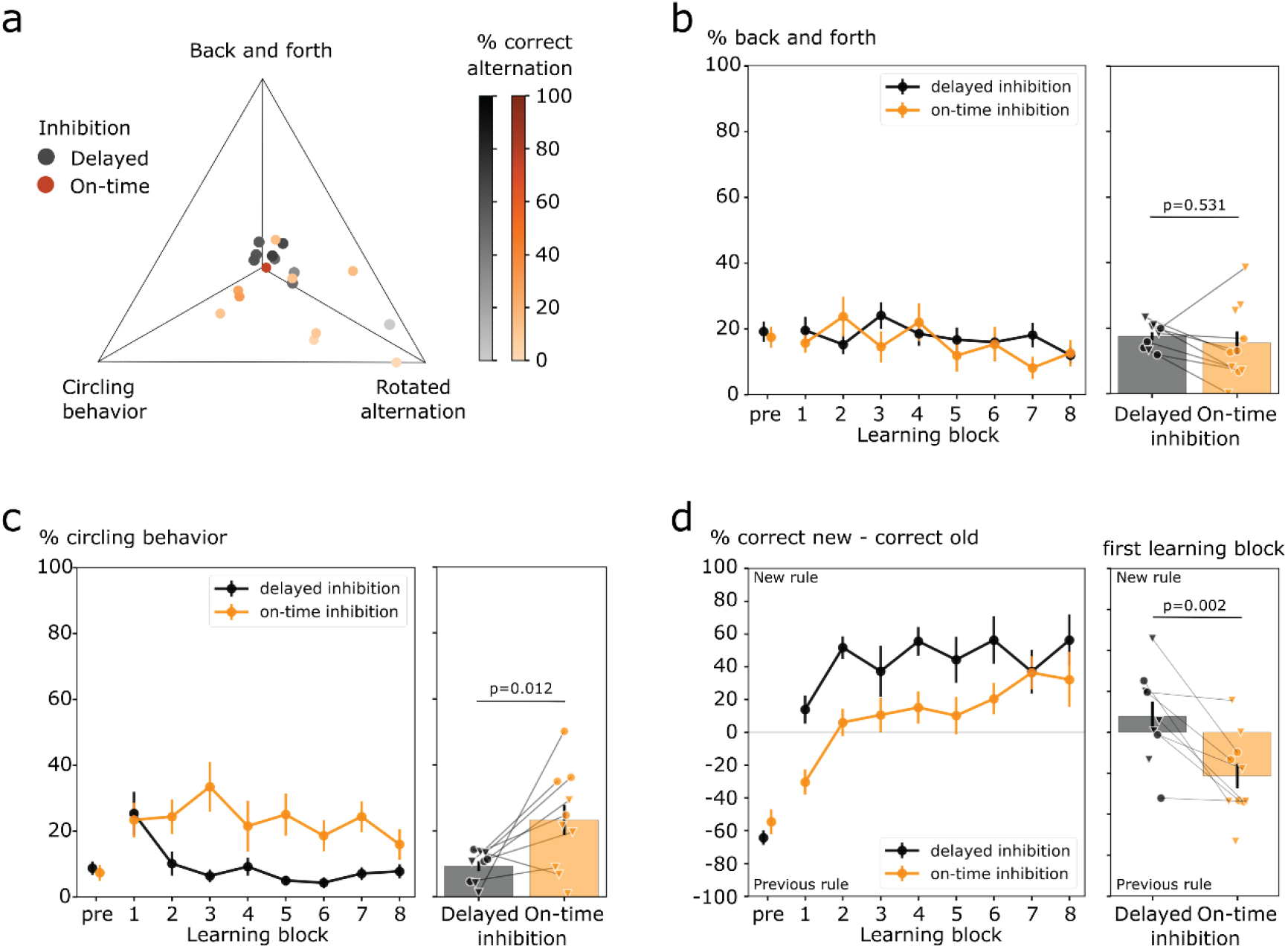
On-time mPFC inhibition leads to perseverative behavior. (**a**) A flattened quaternary plot of the percentages of four types of 4-visit sequences. The dimension representing the percentage correct alternation according to the current rule is color coded. Gray: delayed inhibition, orange: on-time inhibition. (**b**) The percentage of back-and-forth visits was no different between the delayed and on-time inhibition conditions. Left panel: the percentage of back-and-forth visits in each 15-minute block, right panel: the percentage of back-and-forth visits for the whole session. Triangles indicate a rule switch 1 and circles indicate a rule switch 2. (**c**) There was significantly more circling behavior in the on-time inhibition condition compared to the delayed condition. Left panel: the percentage of circular visit sequences in each 15-minute learning block, right panel: the percentage of circular visits for the whole session. Triangles indicate a rule switch 1 and circles indicate a rule switch 2. (**d**) Animals in the on-time inhibition condition continued to perform the previous rule longer than in the delayed condition. Left panel: the percentage of correct alternations according to the previous rule, subtracted from the percentage correct alternation according to the new rule for each 15-minute learning block, right panel: the percentage correct alternation visits new minus previous for the first 15-minute block. Triangles indicate a rule switch 1 and circles indicate a rule switch 2. ‘pre’ indicates the performance on the previous day.

Animals did not engage in circling behavior immediately before a rule switch (Fig. 3c), even though it still leads to a reward in two out of three visits. Following the rule switch, however, animals increased their circling behavior in the first learning block (Fig. 3c). While in the delayed inhibition condition, circling quickly decreased in subsequent learning blocks, animals in the on-time inhibition condition persisted in circling behavior for much longer (Fig. 3c). Across all learning blocks, we observed a significantly higher level of circling behavior in the on-time inhibition condition than the delayed inhibition condition (Fig. 3c; fraction of circling sequences, delayed inhibition: M=0.09, SEM=0.01; on-time inhibition: M=0.23, SEM=0.04; paired samples t-test: t(10)=-3.04, *p*=.012).

Finally, animals may follow one of the two incorrect alternation sequences that only have an expected reward per visit of 0.25. Specifically, after a rule switch, it is expected that animals initially continue to follow the previous rule before having worked out the current rule. To investigate this effect, we looked at the bias towards following the previously learnt rule or the new rule by computing the difference between the percentages of visit sequences that conform to either alternation rule (Fig. 3d). A negative bias indicates a tendency to follow the rule that was in place previously, and a positive bias indicates a switch to the new rule. In the delayed inhibition condition, animals developed a bias to the new rule within the first 15-minute learning block (M=0.09, SEM=0.07), which grew stronger in subsequent learning blocks (Fig. 3d). In the on-time inhibition condition, animals were biased to the old rule in the first learning block (M=-0.27, SEM=0.07) and gradually developed a bias to the correct rule over the course the remaining learning blocks. The rule biases in the first learning block were significantly different between the delayed and on-time inhibition conditions (Fig. 3d; paired samples t-test: t(10)=4.32, *p*=.002).

### Behavioral effects are explained by the timing, not the amount, of optogenetic inhibition

To evaluate the online ripple detections, SWRs were detected offline using an automated algorithm (see Methods) and compared with the online detections. A match rate was defined as the proportion of offline SWRs that were also detected online, and a non-match rate was obtained by taking the proportion of online detected SWRs that did not match any offline detections, see Fig. 4a-b. Compared with the offline algorithm, SWRs were detected with a high sensitivity (match rate on-maze: M=0.79, SEM=0.05, during rest: M=0.80, SEM=0.05), but a lower specificity (non-match rate on-maze: M=0.58, SEM=0.09, during rest: M=0.48, SEM=0.07). Due to an artefact seen in the field potential when the inhibitory LEDs were switched on and off, the match and non-match rates were only calculated for sessions where no inhibition was applied, but the method of online detection was the same for all conditions (see Methods). The online SWR detections occurred primarily at reward locations, and this was no different between the on-time and delayed inhibition conditions (Kolmogorov-Smirnov difference in distributions: K=0.20, *p*=.594; Fig. 4c). Detected SWR rates were higher in the on-time condition than in the delayed condition, both while the animal was on the maze (overall detection rate (Hz), delayed: M=0.30, SEM=0.05; on-time: M=0.53, SEM=0.09; paired samples t-test: t(10)=-2.95, *p*=.014) and in the periods of rest in between learning blocks (overall detection rate (Hz), delayed: M=0.53, SEM=0.04; on-time: M=0.80, SEM=0.10; paired samples t-test: t(10)=-2.98, *p*=.014). Correspondingly, the overall fraction of time that the LEDs were on (total duration of inhibitions divided by total time spent) was higher in the on-time condition compared to the delayed condition (fraction of time light on on-maze, delayed: M=0.06, SEM=0.01; on-time: M=0.09, SEM=0.02; paired samples t-test: t(10)=-2.56, *p*=.028; fraction of time light on rest, delayed: M=0.10, SEM=0.01; on-time: M=0.13, SEM=0.02; paired samples t-test: t(10)=-1.78, p=.106 Fig. 4d). This, however, could not explain the behavioral differences between both conditions, as the fraction of time spent with the inhibitory LEDs on, did not correlate with the alternation performance of the animals (on-maze delayed: Pearson’s r=0.11, *p*=.772; on-time: Pearson’s r=-0.26, *p*=.463; rest delayed: Pearson’s r=0.47, *p*=.205, on-time: Pearson’s r=-0.12, *p*=.732; Fig. 4e). Importantly, sessions within the delayed inhibition condition showed a consistently higher alternation performance, even when the inhibitory LEDs were on for longer. Therefore, the total duration of inhibition itself does not explain the behavioral effects, but rather the timing of the inhibition (directly following a SWR versus with a delay).

**Figure 4.**
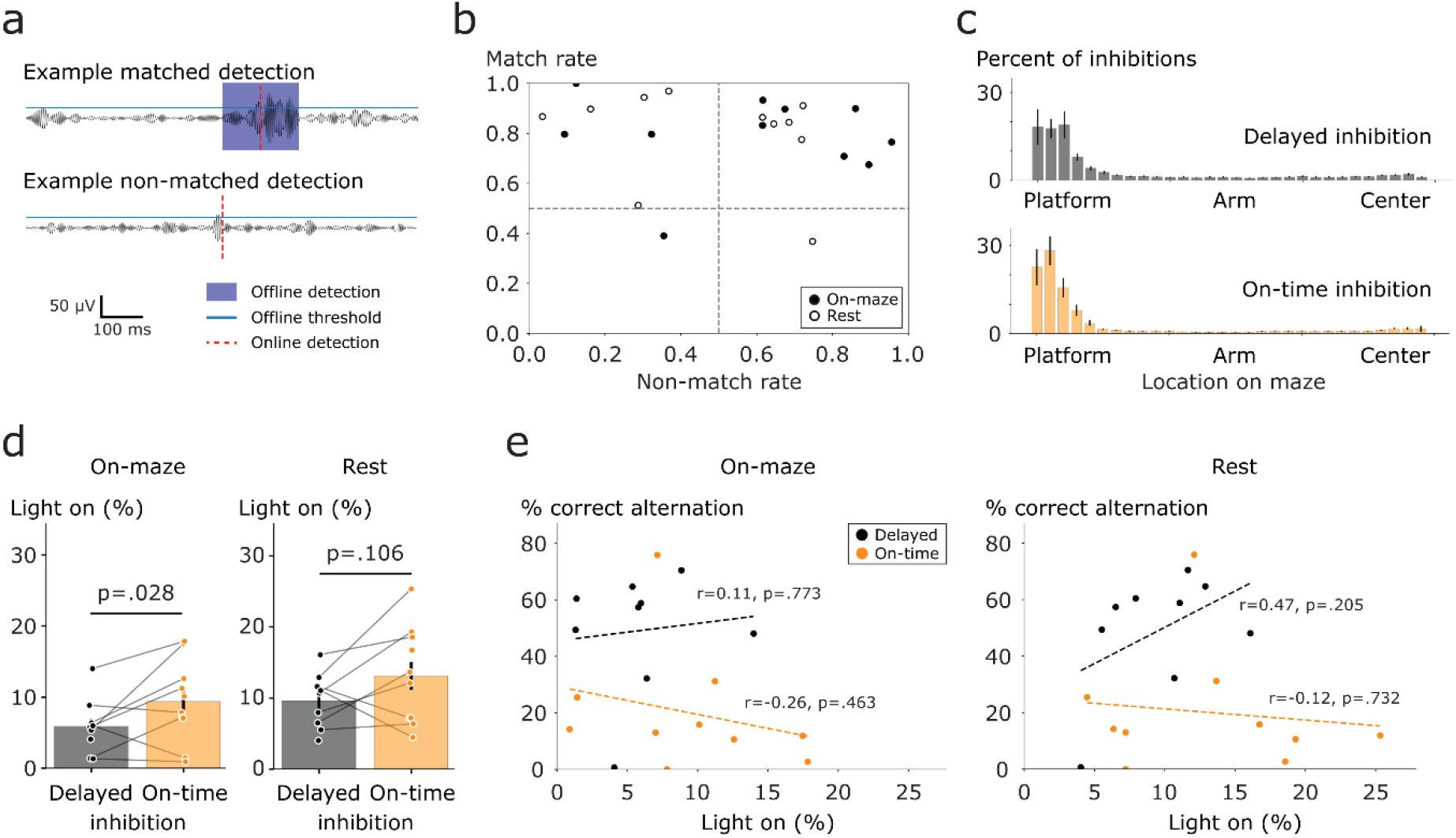
Behavioral effects are explained by the timing, not the amount, of optogenetic inhibition. (**a**) An example of a matched and a non-matched detection, as compared to the offline detection algorithm. Signals are filtered between 160 and 225 Hz. (**b**) The match rate (the number of matched detections divided by the total number of offline detections), and non-match rate (the number of non-matched detections divided by the total number of online detections), for the periods on the maze (solid circles) and the periods of rest in between learning blocks (open circles). (**c**) A histogram of the locations where on the maze the mPFC inhibitions took place during the delayed condition (top panel) and during the on-time condition (bottom panel). (**d**) The percentage of time that the inhibitory LEDs were on, was higher in the on-time condition compared to the delayed inhibition condition. (**e**) The total duration that the inhibitory LEDs were on, did not account for the behavioral differences, as there was no significant correlation between the amount of time the LEDs were on versus the alternation performance while on the maze (left panel) and in the intervening periods of rest (right panel).

## SUPPLEMENTARY INFORMATION

**Supplementary Table S1.** Overview of optogenetic inhibition protocol and spatial alternation rules for all animals in the initial learning and rule switch episodes. Rules 1, 2 and 3 refer to respectively the left, bottom and right arms (from the perspective of the overhead camera) being designated the ‘central’ arm.

| Animal ID | Initial learning |  | Rule switch 1 |  | Rule switch 2 |  |
| --- | --- | --- | --- | --- | --- | --- |
|  | Inhibition | Rule | Inhibition | Rule | Inhibition | Rule |
| B5 | None | 1 | On-time | 3 | Delayed | 2 |
| B9 | None | 2 | Delayed | 1 | On-time | 3 |
| B10 | None | 3 | On-time | 2 | Delayed | 1 |
| B11 | None | 1 | Delayed | 2 |  |  |
| B12 | None | 2 | On-time | 3 |  |  |
| B14 | None | 2 | Delayed | 1 | On-time | 3 |
| B17 | None | 3 | Delayed | 1 | On-time | 2 |
| B18 | None | 1 | On-time | 2 |  |  |
| B20 | None | 3 | On-time | 2 | Delayed | 1 |
| B21 | None | 2 | Delayed | 1 | On-time | 3 |
| B23 | None | 2 | On-time | 3 | Delayed | 1 |

**Supplementary Figure S1.**
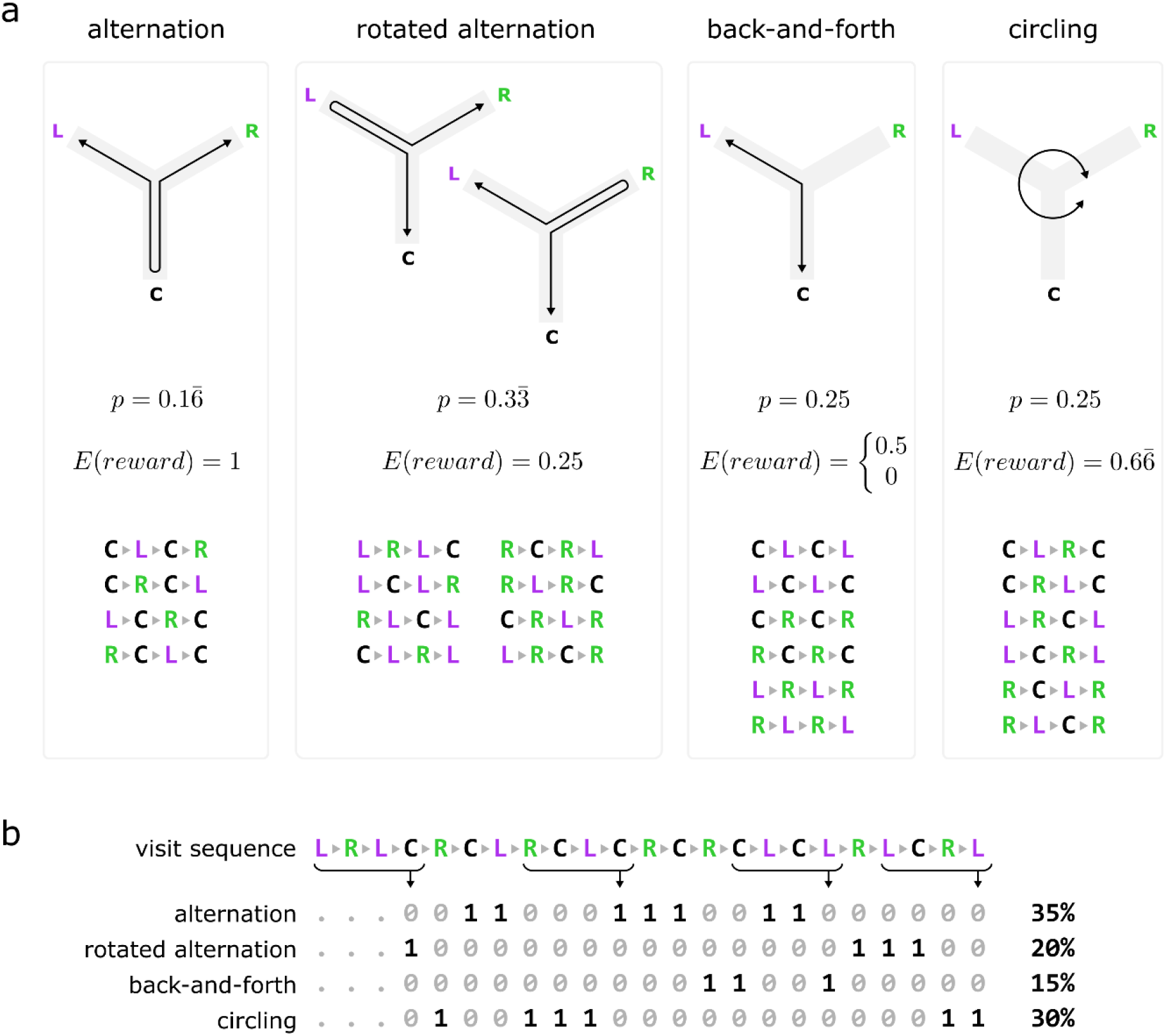
(**a**) Categorization of all possible length-4 visit sequences (without immediate revisits to the same arm) into four groups representing correct alternation behavior, following another (rotated) alternation rule, running back and forth between two arms, and visiting all arms in sequence (circling). For each category, p is the probability that a randomly selected 4-visit sequence is member of this category, and E(reward) is the expected reward per visit. Note that for the back-and-forth category, E(reward) depends on which two arms are part of the pattern. (**b**) Example analysis of a visit sequence in the 3-arm radial maze. Using a sliding window, each length-4 subsequence is categorized according to the groups shown in (a).

## DISCUSSION

In this study, we showed that temporally specific inhibition of mPFC activity in the 200 ms following hippocampal SWRs disrupted the ability of rats to adapt to a rule switch in a spatial alternation task. The performance was compared within-animal to another rule switch in which mPFC activity was inhibited for the same duration (200 ms) but at a delayed interval of 400-450 ms after SWR detection. Our results indicate that the exact timing of the mPFC activity immediately following SWR occurrences in the hippocampus is important for this spatial learning task. This is consistent with previous findings that (1) hippocampal SWRs are important for learning spatial alternation rules^26,27^ and (2) activity in the mPFC is influenced by SWR events in the hippocampus^19,21,31,32^.

According to the two-stage model of memory consolidation, the neocortex provides information to the hippocampus during active exploration while the brain is in a state of slow theta oscillations. The important information then gets reactivated by the hippocampus during fast transient oscillations (SWRs), which in turn provides the neocortex with relevant information necessary for the animal to engage in successful spatial navigation^1^. In our task, the expectation is that during the SWRs, information gets transferred to the mPFC about the location of the reward. The mPFC then takes this feedback and integrates it to form a strategy. In the case of a rule switch, the previously rewarded strategy is no longer rewarding, and the animal must take into account the newly acquired information and adapt its strategy to the changing reward contingencies. We find that with the on-time inhibition, animals perform worse than in the delayed inhibition condition. Therefore, we conclude that this particular time in the interaction is crucial for updating the behavioral strategy when facing changing reward contingencies. Indeed, we see that animals in the delayed inhibition condition can switch to a bias for the new rule within the first 15-minute learning block, while animals in the on-time inhibition condition keep on performing the previously rewarded strategy despite not being rewarded for it anymore.

While our data are consistent with the idea that communication from the hippocampus to the mPFC during SWRs supports learning, there are two alternative explanations. The first is that the mPFC activity may bias the hippocampal network such that reactivation of task-related neural ensembles are favored over reactivation of other ensembles. The biasing would occur separately from the network conditions and precipitating events in the hippocampus that lead to the initiation of SWR bursts. Following this hypothesis, inhibition of mPFC activity during SWRs would decrease task-related replay that supports the learning process. Since the period of mPFC inhibition in our study occurs just past the SWR peak, this hypothesis requires that mPFC would have a continued impact on the expression of hippocampal replay beyond the start of the replay sequence. Future experiments will need to show if mPFC activity indeed impacts the expression of hippocampal replay.

The second alternative explanation is that there is a third brain region or state that influences both the mPFC and the hippocampus simultaneously, which causes SWRs to occur in the hippocampus and influences mPFC activity at the same time. Anatomically, the mPFC and hippocampus are connected through several indirect pathways and one direct pathway^34^. Indirect pathways run through the thalamic nucleus reuniens and through a cortical route, including the lateral entorhinal cortex and the perirhinal cortex. The direct pathway is a monosynaptic projection from the ventral hippocampus to the prelimbic and the infralimbic cortices of the mPFC^34^. If the effect we see is indeed the result of a third brain region, rather than either the mPFC or hippocampus leading the interaction, interesting candidate regions would include the ones in the indirect pathways, for example the nucleus reuniens.

Indeed, it was found that temporarily inactivating the nucleus reuniens with muscimol injections, impaired the acquisition of fear conditioning^35^, but only when the nucleus reuniens was not inactivated during retrieval. When the nucleus reuniens was inactivated during both the acquisition and the retrieval phase, fear conditioning was intact. This indicates that in the absence of the nucleus reuniens, other brain regions may take over, and therefore this region is likely not the (only) structure mediating this interaction.

Previous literature has shown that mPFC inhibition can lead to perseverative behavior^5^, which is referred to as continuously making certain errors, despite the negative (or at least non-positive) feedback. Reversal learning, or rule switching behavior, has also been shown to be particularly impaired following mPFC inhibition^11,12,14^, where animals continue to perform a learnt rule even when it can be deducted that this is no longer the most rewarding strategy. Interestingly, not all animals make the same type of errors. In this dataset, some animals showed a preference for circular trials, and others for a rotated alternation. Animals did not tend to perform a lot of back-and-forth trials, possibly because it was already deducted from the initial learning sessions that all arms need to be visited. It is unclear why certain animals have a tendency for a certain type of error, but in general it is seen that the preference that is there initially, quickly decays in the delayed inhibition condition, where in the on-time inhibition, the animals continue to make the same errors much longer.

As mentioned above, the only direct pathway from the hippocampus to the mPFC is from the ventral hippocampus, and in the current study we are detecting SWRs in the dorsal hippocampus. Importantly, the SWRs in the ventral hippocampus and dorsal hippocampus do not occur at the same times^36^. More specifically, dorsal hippocampal SWRs are well known to occur more during novel and rewarding experiences, whereas ventral SWRs are not modulated by novelty or reward^36^. Furthermore, the timing of dorsal hippocampal SWRs tends to shift with learning, where after the initial learning phase, SWRs occur late in the immobility period, after receiving a reward. In the ventral hippocampus, however, SWRs tend to occur immediately when the animal stops moving, regardless of the stage of learning^36^. As we do see an effect of dorsal hippocampal SWR induced mPFC inhibition, it is likely that the effect we see is through the indirect pathways, rather than the direct pathway from the ventral hippocampus. It would be interesting for future studies to look more at the dissociation between the dorsal and ventral hippocampus and whether the SWR induced mPFC inhibitions lead to similar behaviors when the ventral versus the dorsal hippocampal SWRs are being detected.

Previous studies have attempted to investigate the interaction between the hippocampus and mPFC through contralateral lesions^13,14^. This technique gives some interesting insights but lacks temporal, and some anatomical precision. Another study performed optogenetic inhibition specifically of ventral hippocampal terminals in the mPFC^15^. This technique is anatomically very precise as only those terminals that receive input from the ventral hippocampus, are inactivated. However, in this study mPFC activity was inhibited at task specific times, but not at times specific with respect to neural signatures (i.e., SWRs). An interesting next step would be to target these terminals and inhibit mPFC activity following SWR onset. In this way, both the anatomical and temporal precision would increase, allowing one to draw more precise conclusions about the underlying mechanisms. This experiment would more directly test the importance of the direct monosynaptic pathway from the ventral hippocampus to the mPFC and could be compared with terminal inputs from the indirect pathways, e.g., from the nucleus reuniens.

Some studies have reported differences in SWR modulation in the mPFC depending on whether the animal was asleep or awake^19,32^, and from this one could infer that there may be a difference in the SWR-mPFC mechanisms between the on-maze and resting periods in our task as well. Currently, we detected and inhibited following SWRs during the entire recording session, but in a future experiment it would be interesting to see whether the learning deficits we see are a result of the on-maze SWR-following mPFC activity, or if in fact the SWRs during the periods of rest are playing an important role as well.

Finally, in this study, we implanted optical fibers with an active illumination length of 2.5 mm, allowing to simultaneously inhibit across the different subregions in the mPFC, namely the anterior cingulate, prelimbic, and infralimbic cortices. However, previous research has shown that these regions have distinct functions in the context of fear-related memory^37–39^, cognitive flexibility^33^ and preparatory attention and error-related events^40–44^. An interesting next step for future research would be to inhibit these subregions separately and see whether the behavioral deficits reported here are a result of the inhibition of one specific mPFC subregion in particular or a combination subregion specific activity. As previous studies have pointed at the anterior cingulate cortex in the context of error-related events^40– 44^, one could hypothesize that inhibiting this region leads to a particularly high number of perseverative trials, as the error-related feedback is not processed correctly and therefore the animal is not updating its strategy in the light of the newly acquired information. Another study has suggested that the prelimbic cortex is involved in initiating a new strategy, and the infralimbic cortex helps to establish that new strategy^33^, meaning that the timing of the inhibitions (early after the rule switch or later in the learning session) may involve different mPFC subregions. These hypotheses have yet to be tested in a closed-loop SWR induced manipulation paradigm.

In conclusion, our results show that the specific timing of hippocampal SWRs, related to the mPFC activity, seems to be a key element that drives behavior in this type of spatial learning. Inhibiting the mPFC activity at the precise time of SWR onset, led to a decrease in behavioral performance and an increase in perseverative behavior and caused animals to be less able to switch to a new strategy in the light of changing reward contingencies.

## METHODS

### Animals

Adult male Long Evans rats (N=11; 300-400 g at implantation) were used in this study. The animals were kept on a 12-hour light/dark cycle. Upon arrival, the rats were housed in pairs and had *ad libitum* access to food and water. Throughout the experiment, animals were exposed to an enriched environment (two-level play pen) for about one hour per day during which they could interact with their cage mate, run on a wheel and forage for food.

### Apparatus

Experiments were performed in a 2.9 by 4.2 meters room with black walls. A three-arm radial maze, elevated 50 cm above the ground, consisted of three equal-length arms (62 cm) connected to a 38 cm by 38 cm central platform with 120-degree angle between arms. Each arm ended in a 22 cm by 22 cm platform with a small liquid reward well that was connected to a peristaltic pump. Access from the central platform to each arm was controlled by three automated doors, which opened at the beginning of each learning block and closed at the end. The food pumps and doors were controlled by software over Wi-Fi and reward was automatically delivered by live tracking of the animals’ location and responses in the maze using software developed in-house (Hive, https://bitbucket.org/kloostermannerflab/fklab-controller-lab).

### Pre-training

Prior to experimental sessions, rats were familiarized to the experimental room and pre-trained to run for reward (sweet milk) on an elevated 148 cm-long linear track. A 22 cm × 22 cm platform with liquid reward pump was located at both ends of the track. Each training day consisted of two 15-minute blocks separated by a 15-minute break in a dark resting box. The animals had to run back and forth and received a food reward at each end of the track. The pre-training was completed once animals reached the criterium of more than 100 rewarded platform visits in the two 15-minute blocks combined. Animals took 8 to 13 days to complete pre-training.

### Continuous spatial alternation task

The goal of the behavioral task was for the rats to learn one of three possible spatial alternation rules in the three-arm radial maze. For a given rule, one arm was designated “center” arm and animals had to learn to alternately visit the two other arms (referred to as “left” (from center) and “right” (from center) arms) using the center arm as a start and end point. There were no physical cues to indicate the identity of the center, left and right arms, so animals had to deduce the current rule through trial and error. For each visit to an arm that was consistent with the alternation rule, animals received a reward (sweet milk). Learning episodes were split into up to eight 15-minute learning blocks per day, separated by 15-minute breaks in a dark resting box. The first day of the initial learning and rule switching episodes always consisted of eight learning blocks, which was followed by two extra days where animals were trained until reaching the learning criterium of 80% correct overall (minimum 2 and maximum 8 learning blocks). The order in which the three rules were presented was counterbalanced across animals (see Supplementary Table S1).

### Surgical procedures

To record electrophysiological signals from the hippocampus and inhibit mPFC activity while the animals were freely moving on the maze, a 3D-printed micro-drive array was designed and fabricated (printed with Grey resin, Form 3+ printer, Formlabs, Somerville, MA, USA) to house eight tetrodes targeting the hippocampus and two tapered optical fibers targeting the mPFC (tapered lambda fiber cannulas, 0.66 Numerical Aperture, 2.5 mm active length, 20 mm implant length, Optogenix, Arnesano, Italy) rigidly fixed at a distance of 0.9 mm. The tapered optical fibers had an active (illumination) length of 2.5 mm, allowing simultaneous inhibition of the anterior cingulate, the prelimbic and the infralimbic cortex of the mPFC. The optical fibers were connected to customized LEDs (angled LEDs 595nm, Doric Lenses, Québec, Canada) with fiber-optic cannula assemblies.

Animals were anesthetized in an induction chamber with 5% isoflurane and throughout the surgery anesthesia was maintained via a nose cap. Vital signs were monitored continuously (PhysioSuite monitoring system, Kent Scientific, Torrington, CT, USA) and the level of isoflurane (1-2%) was adjusted to maintain a constant level of anesthesia. A heating pad with a rectal probe feedback loop kept the body temperature constant. The animal’s head was placed in the stereotaxic frame using ear bars for stabilization, and eye ointment and aluminum foil to protect the eyes. The head was shaved, and the skin was thoroughly disinfected before making an incision that ranged from anterior to the bregma to posterior to lambda. The skull was exposed using a retractor and then thoroughly cleaned and scored with a scalpel to improve adhesion with dental cement at the end of the procedure. Craniotomies were drilled above the right hippocampus (2 mm diameter, centered 3.4 mm posterior to Bregma and 2 mm from the midline) and bilaterally above the mPFC (2.7 mm anterior to Bregma, extending 1 mm on both sides of the midline). For optogenetic inhibition of the mPFC, the viral vector pAAV-CamKII-ArchT-GFP (titer: 1.9×10^13^ genome copies per ml, Addgene, Watertown, MA, USA) was injected bilaterally 0.45 mm from the midline at 1, 2, 3 and 4 mm depth from the brain surface, using 0.5 μl per injection, with a rate of 0.6 μl per minute and 5 minutes rest after each injection. Injections were performed using a stereotaxic injector (quintessential stereotaxic injector, Stoelting company, Wood Dale, IL, USA) and a glass pipette with a tip diameter of 30 μm attached to a Hamilton syringe.

To anchor the micro-drive array, 8-10 burr holes were drilled around the perimeter of the skull and small bone screws (1.19 × 4.8 mm self-tapping bone screws, 19010-00, Fine Science Tools, Foster City, CA, USA) were carefully inserted. One screw was used as electrical ground. Light-curable dental cement (wave A2 syringes, Dental Elite, Lille, France) was used to fix the micro-drive array in place. Before the animal was woken up, the skin was disinfected once again, and a topical anesthetic was applied on the wound (lidocaine cream). At this point, and in the three days after, 0.35 ml Metacam was injected subcutaneously as an analgesic. The wound healing was monitored every day and antibacterial cream (Picri-Baume) was applied when necessary. After surgery, animals spent two weeks in quarantine and were single housed. Once fully recovered from surgery, animals were kept at a mild food restriction schedule, maintaining 85-95% of their original body weight. The two optical fibers were lowered to their final depth of 4 mm during surgery. All hippocampal tetrodes could be moved independently and were lowered slowly into area CA1 in the weeks after surgery.

### Online SWR detection and mPFC inhibition

Most tetrodes were positioned near the pyramidal cell layer of area CA1, using feedback from electrophysiological recordings to optimize the amplitude of ripple oscillations. One tetrode was used as a reference and positioned in the white matter overlying the hippocampus, and one tetrode was left in the cortex. Neural signals (4 kHz) and overhead video images (50 Hz) were acquired with a Digilynx 16SX acquisition system and Cheetah software (Neuralynx, Bozeman, MT, USA).

Electrodes from one to three tetrodes with strong ripple activity and high signal-to-noise ratio were manually selected for online detection of hippocampal SWR events. Digitized signals were streamed to a dedicated workstation running Falcon software^45^. The streamed signals were filtered between 130 and 283 Hz using a Chebyshev type-II IIR filter, and an envelope was estimated using the squared value of the filtered signals. A running threshold was used to detect putative SWR events. Thresholds were based on sum of the computed envelopes, and obtained by taking the mean of the summed signal, plus N times the mean absolute deviation, where the multiplier N was set manually based on the signal-to-noise ratio of the signal and ranged between 12 and 16. Potential false positive detections were reduced by applying the same filter to a signal recorded by an electrode in the cortex and discarding hippocampal SWR detections that were also detected in the cortex (within a window of -40 ms to 1.5 ms from the hippocampal detection). The threshold for the cortex detection was calculated in the same way as in the hippocampus and the multiplier was set to 12. Each hippocampal SWR detection was logged in the Cheetah software through a TTL pulse from an Arduino UNO to the Digilynx acquisition system. A 50 ms lock-out period was applied after each detection to avoid continuous detections of one single event. On the first day of a rule switch episode, each hippocampal SWR detection was followed by activation of the two LEDs (200 ms, 200 mA) to inhibit mPFC activity either immediately (on-time inhibition condition), or after a randomized delay of 400-450 ms (delayed inhibition condition). The activation of the two LEDs was driven via a LED controller (SLC-AV04-US, Mightex, Toronto, Canada) through the tether of the Digilynx acquisition system. Switching the LEDs on and off caused an artefact in the field potential signal, and therefore another 50 ms lock-out period was applied for each time the LEDs were turned on and off. The temporal order (i.e., rule switch 1 or 2) of the on-time and delayed conditions was counterbalanced across animals. Hippocampal SWR events were detected online both during learning blocks and intervening rest periods.

### Histology

At the end of an experiment, animals were euthanized with an intraperitoneal injection of 5 ml pentobarbital (Dolethal, 200 mg/ml) and electrolytic lesions (60 µA for 10 seconds on each electrode) were made to mark the location of the electrode tips in the hippocampus. Following transcardial perfusion with 4% formaldehyde, brains were stored in a tube with 4% formaldehyde in a cold room (4°C) for 24 hours. After this, brains were transferred to a tube with 30% sucrose solution for several days until they sank to the bottom (three days on average). The brains were next sliced at 100 µm thickness using a vibratome (VT1000 S, Leica, Wetzlar, Germany) and left to dry overnight on microscope slides. Brain sections that contained the dorsal hippocampus were stained with a non-fluorescent cresyl violet Nissl stain to visualize the cell bodies and determine the position of the electrodes. Brain sections that contained the mPFC were stained with a fluorescent Nissl stain (NeuroTrace 530/615 Red Fluorescent Nissl Stain, Thermo Fisher, Waltham, MA, USA), and expression of the viral construct was evaluated by visualizing GFP with a fluorescent microscope (MVX10, Olympus, Tokyo, Japan).

### Data analysis

The position of the animals was tracked by training a deep neural network (DeepLabCut^46^) to identify the tail base and two colored LEDs attached to the implant in the overhead video data. Training was performed once on 407 video frames from 68 recording sessions in 10 animals, for 500,000 iterations. The trained network was then used to track the animals’ position in all video frames. Post-processing was done using in-house developed software and included removing jumps of more than 20 pixels and smoothing the velocity with a gaussian kernel (bandwidth = 250 ms). Based on the obtained position data, sequences of visits to the reward platforms were then detected and used to analyze task performance and behavioral patterns.

To determine the quality of online SWR detection, SWR events were detected offline post-hoc and compared with the online detected SWRs. Offline detection was performed by applying a band-pass filter to the signals (160-225 Hz) and computing the envelope as the absolute value of the Hilbert transform. High and low thresholds were set respectively for detection of SWR events and determining the start and the end times of these events. The thresholds were obtained by using the median amplitude of the signals plus N times the median absolute deviation. The multiplier N was set between 6 and 9 (high threshold; for detection) and to 1 for the low threshold (for determining the start and end times). A non-match rate was defined as the fraction of online detected SWRs that were not classified as a SWR with the offline algorithm, and the match rate was defined as the fraction of offline detected ripples that were also detected online. As LEDs for optogenetic inhibition were placed on the implant close to the recording electrodes and headstage, switching on and off the LEDs caused an electrical artefact in the recorded field potential in the hippocampus. For this reason, the match and non-match rates were only calculated for the initial learning sessions, in which no artefacts were present and online SWR detection was performed the same as for delayed and on-time inhibition conditions.

### Statistics

The difference in task performance between the on-time and delayed inhibition conditions was evaluated using *t*-tests from the SciPy library for python^47^. As some animals were not able to participate in both conditions due to technical issues with the implant (see Supplementary Table S1), we report paired samples t-test where the missing sessions (N=3) were imputed using the median of all other sessions within that condition.

## Acknowledgements

We thank Lies Deceuninck for her contributions in conceptualization of the behavioral paradigm and helping with troubleshooting during experiments, Marine Guyot for her software contributions, Louise Vanmarcke for her help in optimizing the virus injections, Meriam Malekzadeh for her help in testing the behavioral paradigm and James Coleman for his help in designing a prototype of the micro-drive array. F.K. and J.-J.S. are funded by Research Foundation Flanders (FWO), Belgium under grant number G0A5422N. F.K. is funded by Research Foundation Flanders (FWO), Belgium under grant number G077321N and KU Leuven, Belgium C1 grant C14/17/042.

## Author contributions

Conceptualization, J.-J.S., H.d.B. and F.K.; Methodology, J.-J.S., H.d.B. and F.K.; Software, F.K.; Formal analysis, H.d.B, M.V.D and F.K.; Investigation, H.d.B., M.V.D. and J.-J.S.; Writing - Original draft, H.d.B, M.V.D. and F.K.; Writing - review and editing, H.d.B., M.V.D., J.-J.S. and F.K.; Visualization, H.d.B. and F.K.; Supervision, F.K.; Funding Acquisition, F.K. and J.-J.S.

## Materials & Correspondence

Correspondence and request for materials can be addressed to Fabian Kloosterman.

## Data Availability

The data that support the findings of this study are available from the corresponding author upon reasonable request.

